# Pioneer factor Foxa2 mediates chromatin conformation changes in ligand-dependent activation of nuclear receptor FXR

**DOI:** 10.1101/2023.03.06.531297

**Authors:** Yi Hao, Lu Han, Anqi Wu, Irina M. Bochkis

## Abstract

Activation of nuclear receptors, a family of ligand-dependent transcription factors, is used extensively in development of drug targets. We have previously shown that pioneer factor Foxa2 opens chromatin for binding of nuclear receptors FXR and LXRα during acute ligand activation. FXR is activated by bile acids and deletion of Foxa2 in the liver results in intrahepatic cholestasis. We hypothesized that Foxa2 also enables chromatin conformational changes during ligand activation. We performed Foxa2 HiChIP to assess Foxa2-dependent long-range interactions in mouse livers treated with either vehicle control or FXR agonist GW4064. HiChIP contact analysis shows that global chromatin interactions are dramatically increased during FXR activation. Ligand-treated livers exhibit extensive redistribution of topological associated domains (TAD and substantial increase in Foxa2-anchored loops, suggesting Foxa2 is involved in dynamic chromatin conformational changes. We demonstrate that chromatin conformation, including genome-wide interactions, TADs, intra-chromosomal and inter-chromosomal Foxa2-anchored loops, drastically changes upon addition of FXR agonist. Hence, we determine a novel role for Foxa2 in enabling these conformational changes, extending its function in bile acid metabolism.

## Introduction

Activation of nuclear receptors, a family of ligand-dependent transcription factors, is used extensively in pharmacology to develop drug targets for diverse medical conditions, including metabolic disease and cancer ^1-4^. Farnesoid X receptors (FXR), liver X receptors (LXR), and peroxisome proliferator-activated receptors (PPAR), members of the nuclear receptor family, are essential for metabolic homeostasis, regulating targets in bile acid, cholesterol, fatty acid, and glucose metabolism ^5^. A previous model regarding ligand activation of these receptors involved a ligand-independent binding mechanism, depending on co-repressor/co-activator exchange leading to activation of gene expression with addition of the ligand. However, we have shown that FXR and LXRα exhibit both ligand-independent and ligand-dependent binding, with chromatin accessibility being induced during ligand activation allowing for additional binding. These changes and activation of ligand-responsive gene expression require pioneer factor Foxa2 ^6^.

We have characterized a comprehensive role for winged-helix transcription factor Foxa2 in hepatic bile acid metabolism. Deletion of Foxa2 in the liver leads to intrahepatic cholestasis, and expression of FOXA2 is markedly decreased in liver samples from individuals with different cholestatic syndromes^7^. In addition, bile acid-dependent activation of nuclear receptor FXR to induce gene expression to protect liver from injury requires Foxa2 ^8^. Furthermore, we demonstrated that that bile acid-induced inflammation in young *Foxa2* mutants, once chronic, affected global metabolic homeostasis as they aged, leading to age-onset obesity ^9^.

In this study, we hypothesized that Foxa2 enables chromatin conformational changes during ligand activation of FXR. We performed Foxa2 HiChIP to assess Foxa2-dependent long-range interactions in mouse livers treated with either vehicle control or FXR agonist GW4064. HiChIP contact analysis shows that global chromatin interactions are substantially increased during FXR activation. We demonstrate that chromatin conformation, including genome-wide interactions, TADs, intra-chromosomal and inter-chromosomal loops, drastically changes upon addition of FXR ligand. Hence, we determine a novel role for Foxa2 in enabling these conformational changes, extending its function in bile acid metabolism.

## RESULTS

### Global chromatin interactions are increased during FXR activation

We demonstrated previously that Foxa2 plays an extensive role in bile acid metabolism ^7-9^ and is required for ligand-dependent activation of FXR ^6^. To test the hypothesis that, in addition to opening chromatin, Foxa2 is involved in chromatin conformation changes to enable ligand-dependent activation of FXR, we performed Foxa2 HiChIP (Hi-C & ChIP, study design in Fig. 1, HiChIP valid pairs metrics in Extended Data Fig. 1) in mice treated with either vehicle control or FXR ligand GW4064. HiChIP paired-end reads were mapped and filtered for valid interactions using HiC-Pro ^10^. Hi-C maps of all valid interactions show more chromatin loops in ligand-activated condition than in vehicle control (Fig. 2a). Next, genome wide intra-chromosomal bin pairs were filtered for an interaction distance between 20kb and 2Mb to identify statistically significant interactions. Representative arc-plots of significant chromatin interactions in both conditions on chromosome X are shown in Fig. 2b, with more observed in ligand-activated livers. Number of total significant interactions increases with addition of FXR agonist (14778 total interactions specifically for vehicle, 27357 specifically for GW4064, 93480 common for both, bin size 50kb, FDR < 0.001, Fig. 2c). Scanning motif analysis of positional weight matrices in the JASPAR and TRANSFAC databases in genomic regions associated with significant interactions identified highly enriched forkhead motif bound by Fox transcription factors in both conditions (p-value ∽0), as expected for Foxa2 ChIP. Then, we performed differential significant contact analysis between two conditions (statistics per chromosome in Extended Data Fig. 2a, 31,891 total). An example of a genomic region (chr11: 17600000 -17800000) with increased chromatin interactions in ligand-treated condition is shown in Fig. 2d. A virtual 4C plot shows increased signal in GW-treated livers and the difference (delta) between two conditions (Fig. 2e). To identify functional differences in differential interactions we mapped these regions to closest genes using GREAT^11^ and performed Ingenuity Pathway Analysis (IPA). Overrepresented pathways included “Oxidative Stress”, “Fatty Acid Metabolism” and “Acute Phase Response” for genes in interaction regions in control livers while genes in FXR and CAR activation pathways and those regulating cholesterol metabolism were found in genes in interaction regions in agonist-treated livers, consistent with bile acid activation (Extended Data Fig. 2a). IPA of genes in differential interaction regions identified multiple upstream regulators, including HNF4α, CEBPB, and EP300 for control livers and NFE2L2, SMARCB1, and chenodeoxycholic acid (Extended Data Fig. 2b) as well chenodeoxycholic acid-regulated network (Extended Data Fig. 2c) for GW-treated livers. Chenodeoxycholic acid is a naturally occurring bile acid that activates FXR. Hence, IPA analysis confirms FXR activation in agonist-treated livers.

**Figure 1.**
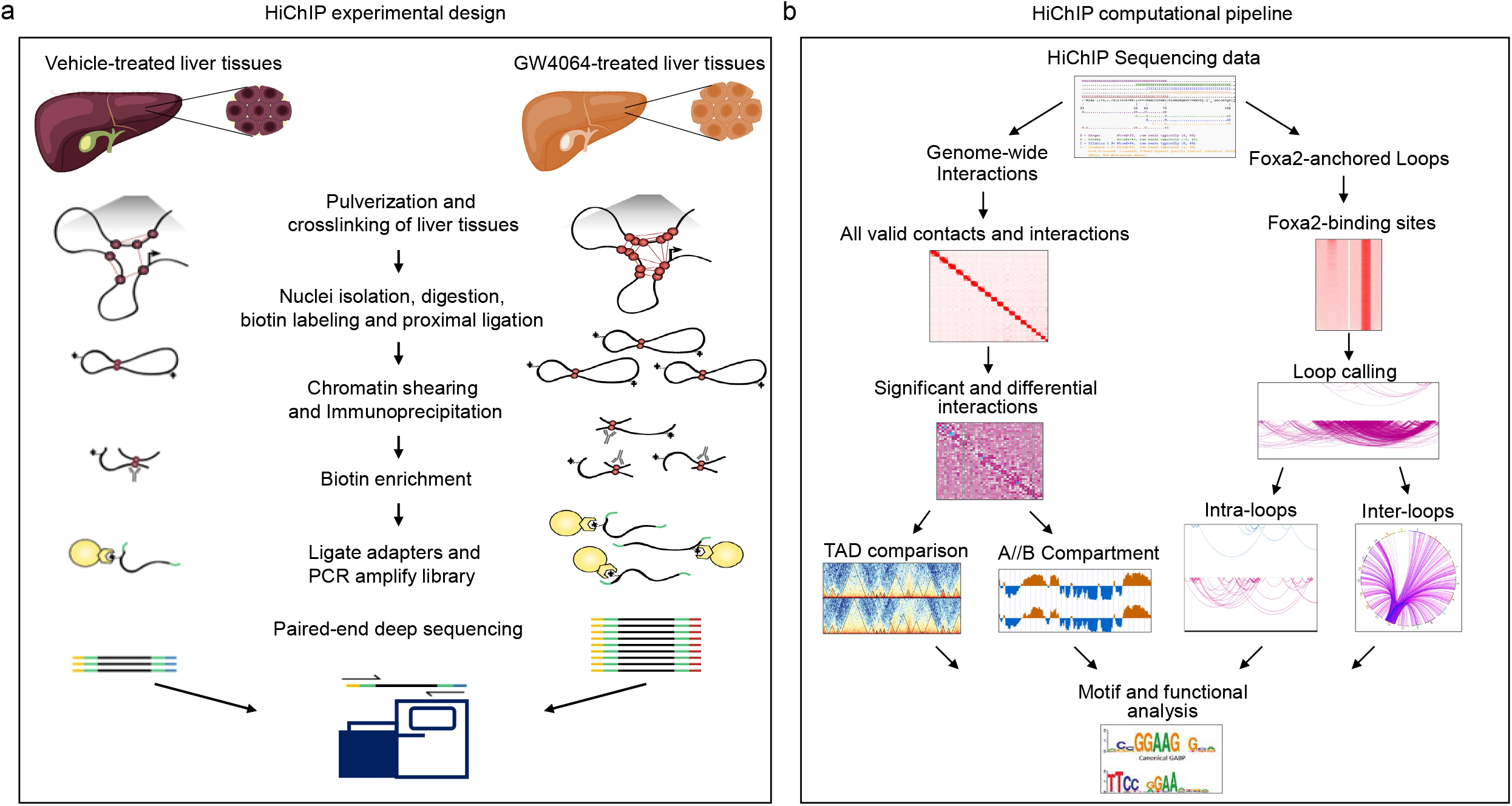
Study design and computational analysis pipeline. (**a**) HiChIP experimental design. Liver tissues were pulverized and crosslinked. Nuclei were isolated, nuclear membranes were permeabilized, chromosomes were digested with both MboI and HinfI, and the ends of DNA fragments were biotin-labeled. The proximal ends were ligated. Then crosslinked chromatin was sheared into fragments. Foxa2 antibody was added and Foxa2-bound chromatins were immunoprecipitated. The HiChIP DNA fragments were enriched for biotin and were ligated with adapters. The HiChIP libraries were amplified by PCR and sent for paired-end deep sequencing. (**b**) HiChIP computational pipeline. First, we identified genome-wide valid interactions, significant interactions, and differential interactions in both vehicle control and GW4064-treated livers. We proceeded to call topological associating domains (TADs) and compartments (activated Compartment A and repressed Compartment B) form genome-wide valid interaction data. In parallel, we used peaks from Foxa2 ChIP-Seq ^6^ that also have signal in HiChIP data to identify Foxa2-anchored chromatin loops, divided into intrachromosomal and interchromosomal. We performed scanning motif analysis of sequences in these regions and mapped them to genes, which were used subsequently for functional analysis.

**Figure 2.**
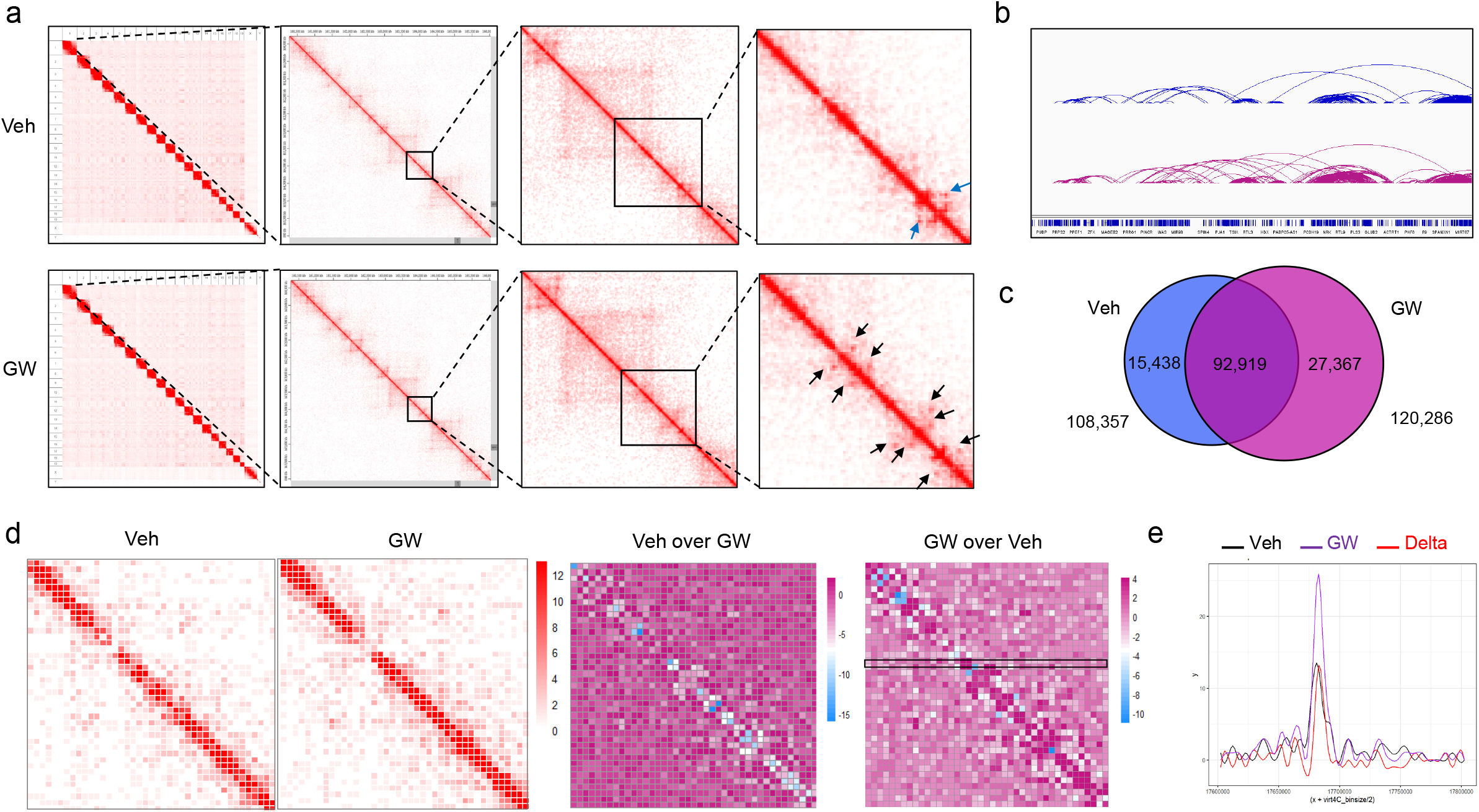
Genome-wide chromatin interactions increase during ligand-dependent activation of FXR. **(a)** Hi-C maps of all valid interactions in vehicle and GW4064-treated livers visualized with Juicebox ^17^ at increasing zoom scales from left to right. Red squares along the diagonal in the left-most panel (top for vehicle, bottom for GW4064) represent the interactions within each chromosome territory. In the middle two panels (top for vehicle, bottom for GW4064), TADs are shown in plaid pattern along the diagonal. In the right-most panel (top for vehicle, bottom for GW4064), the arrows indicate presence of “corner peaks” of chromatin interactions at the edges of TADs, revealing the presence of chromatin loops. The blue arrows indicate interaction loops unique to vehicle treatment, whereas the black arrows specify those specific to GW4064 treatment, showing more chromatin loops with addition of agonist. (**b**) Integrative Genome Viewer (IGV) ^18^ view of representative arc-plots of significant chromatin interactions in vehicle- and GW-treated conditions on chromosome X. (**c**) Venn diagram showing that the numbers of significant chromatin interactions increased with the addition of FXR agonist (14778 total interactions specifically for vehicle, 27357 specifically for GW4064, 93480 common for both, under bin size of 50kb and with the FDR values < 0.001). (**d**) Comparison of differential interactions within a specific genomic region (chr11: 17600000 -17800000), showing increased chromatin interactions with GW4064 treatment compared to vehicle in this region. Two left-most panels display the Hi-C maps along the diagonal within this genomic region for vehicle and GW4064 treatment, respectively. The third panel illustrates differential interactions comparing vehicle to GW4064 HiChIP signal along the diagonal, in white and blue colors, corresponding to decreased interactions in vehicle treated livers. The fourth panel shows differential interactions comparing GW4064 to vehicle HiChIP signal along the diagonal, in dark purple color, corresponding to increased interactions in GW4064. (**e**) Virtual 4C plot comparing chromatin interactions from a single genomic location (chr11: 1768000) to the rest of the genomic region (chr11: 17600000-17800000), corresponding to the region in black rectangle in the right-most panel in Figure 4d.

### FXR agonist treatment significantly changes TAD distribution

We proceeded to call topological associating domains (TAD) in our HiChIP data and found a comparable number in vehicle and GW-treated conditions (3575 for vehicle, 3734 for GW 4064). However, only a third of the TADs overlapped, while majority of TADs differed (1295 overlapping TADs, 2436 differential TADs, Fig. 3a). Distribution of differential TADs on each chromosome is shown in Extended Data Fig. 3a. Next, we mapped differential TADs to closest genes using GREAT^11^ and selected 1000 genes that mapped closest to the transcription start site (TSS) for pathway analysis with Enrichr ^12^. Overrepresented pathways included “LXR regulation of gluconeogenesis” and “FXR and LXR regulation of cholesterol metabolism”, consistent with bile acid activation, as well as “Cohesin loading onto chromatin”, congruous with cohesin function in mediating chromatin loop formation (Extended Data Fig. 3b, top panel) ^13^. We also utilized published ChIP-Seq data sets in the mouse genome and intersected differential TAD regions and the binding data present in the ChEA database, identifying overlap with RUNX1, TP53, β-catenin, and LXR targets (Extended Data Fig. 3b, bottom panel, ChEA analysis in Enrich). RUNX1 improves bile-acid induced hepatic inflammation in cholestasis ^14^, p53 plays a role in bile acid metabolism by regulating SHP (*Nr0b2*) ^15^, and β-catenin regulates FXR-dependent gene expression in cholestasis ^16^. Hence our analysis of differential TADs identified regulators important for bile acid homeostasis and shows that ligand-dependent activation of FXR is associated with substantial changes in TAD distribution.

**Figure 3.**
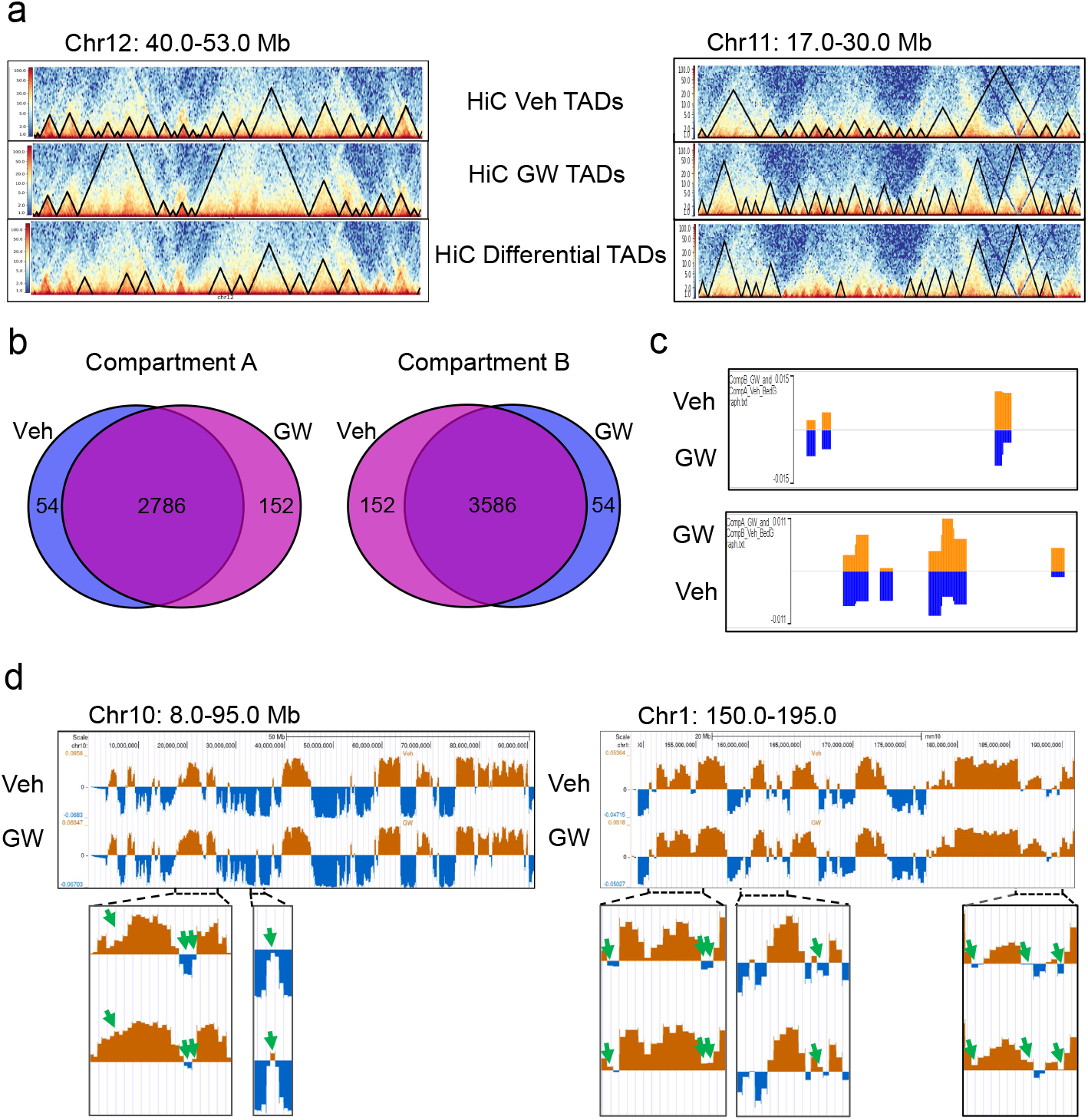
Topological associating domain (TADs) and A/B compartments are reshuffled with the addition of FXR ligands. (**a)** Comparison of TAD distributions in vehicle and GW4064 for chromatin regions chr12: 40-53.0 Mb (left panels) and chr11: 8.0-20.0Mb (right panels). TADs in vehicle control (top panels), TADs with agonist treatment (middle panels), differential TADs (bottom panels). The black triangles in each panel indicate the separated TAD domains in each condition. (**b**) Venn diagram comparing activated and repressed regions (Compartment A and Compartment B, respectively) in HiChIP interactions showing that 54 such regions changed from being activated in vehicle-treated to repressed in GW-treated livers while 152 moved from Compartment A in agonist-treated to Compartment B in control livers. (**c**) Examples of compartment switching (Compartment A in vehicle control to Compartment B in GW4064 treatment, top panel; Compartment B in vehicle control to Compartment A in GW4064 treatment, bottom panel). Orange color indicates compartment A (active), while blue color indicates compartment B (inactive). (**d**) Representative comparison of A/B compartments in vehicle control and GW4064-treated livers (chr10: 8.0-95.0Mb, left; chr1: 150.0-195.0Mb, right). Orange color indicates compartment A (active), while blue color indicates compartment B (inactive). The green arrows point at regions switching from inactive in vehicle control to more active in GW4064-treated livers.

Next, we identified activated and repressed regions (Compartment A and Compartment B, respectively) in HiChIP interactions and found that 54 such regions changed from being activated in vehicle-treated to repressed in GW-treated livers while 152 moved from Compartment A in agonist-treated to Compartment B in control livers (Fig. 3b). Examples of full and partial switching are shown in Fig. 3c, 3d. Functional analysis of genes in these regions shows “Cytoskeletal Organization”, “Reactive Oxygen Species”, and “RXR/VDR pathway” pathways overrepresented in 54 regions repressed with agonist treatment and “Proteins with Altered Expression in Aging”, “Sodium/Proton Exchangers”, and “Opening of Calcium Channels” in 152 regions activated with GW treatment (Extended Data Fig. 3c).

### Foxa2-anchored loops are drastically increased genome-wide and at bile acid targets

To relate Foxa2 binding sites to changes in chromatin conformation, we used previously called bound regions in Foxa2 ChIP-Seq (7306vehicle, 22666 GW4064) ^6^ and found that they are also occupied by Foxa2 in HiChIP data, as expected (plot, Fig. 4a, heatmap, Fig. 4b). Scanning motif analysis of positional weight matrices in the JASPAR and TRANSFAC databases in Foxa2 binding sites identified a highly enriched forkhead motif bound by Fox transcription factors in both conditions (p-value ∽0), as expected for Foxa2 ChIP. We used these regions to identify Foxa2-anchored intrachromosomal loops (3119 for vehicle, 18503 for GW4064, Fig. 4c, HiChIP valid pairs metrics in Extended Data Fig. 4). Although 70 percent of sites bound by Foxa2 in vehicle-treated livers overlapped with regions occupied in livers activated by GW4064 ^6^, only about a third of intrachromosomal loops were common to both conditions. A representative region showing more Foxa2-anchored intrachromosomal loops with agonist treatment is shown in Fig. 4d.

**Figure 4.**
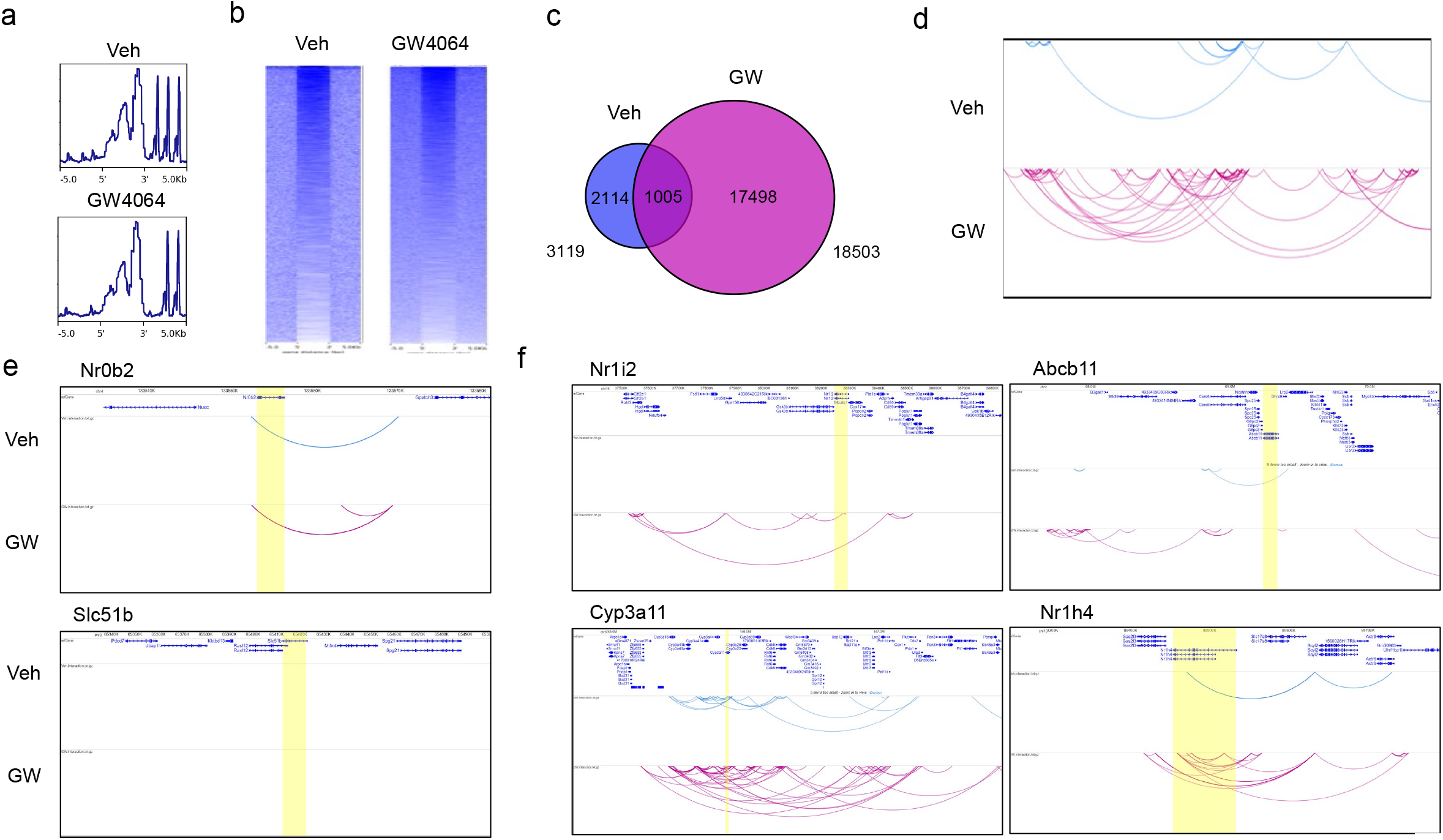
Foxa2-anchored intrachromosomal loops are drastically increased during ligand-dependent activation of FXR. **(a**) Foxa2 HiChIP signal (RPKM) plot and (**b**) heatmaps at Foxa2 binding sites from Foxa2 ChIP-Seq (7306 sites vehicle, 22666 GW4064) ^6^ (**c**) Venn diagram comparing Foxa2-anchored intrachromosomal loops in vehicle and GW4064-treated livers. There are totally 3119 intra-loops unique to vehicle control, 18503 intra-loops unique to GW4064 treatment, and 1005 intra-loops common to both conditions. (**d**) Representative arc plot showing total Foxa2-anchored intra-chromosomal loops within the genomic region chr10: 66.7-69.8 Mb in vehicle and GW4064 treatment. (**e**) Representative arc plots showing intrachromosomal loops at FXR targets with ligand-independent binding (*Nr0b2 & Slc51b*). (**f**) Representative arc plots showing the number of Foxa2-anchored intrachromosomal loops drastically increases at a number of FXR targets with ligand-dependent FXR binding, including its own locus (*Nr1i2*/PXR, *Abcb11*/Bsep, Cyp3a11, *Nr1h4*/FXR)

We have demonstrated that contrary to the established paradigm, FXR binding changes drastically with addition of an agonist and ligand-dependent FXR binding is Foxa2-dependent ^6^. We proceeded to ascertain whether Foxa2-anchored intrachromosomal loops differed at FXR ligand-independent and ligand-dependent sites. FXR binding is ligand-independent at *Nr0b2* locus ^6^ and FXR target *Slc51b* is expressed in the intestine but not in the liver. There are no substantial changes in loops at *Nr0b2* locus and we observe no Foxa2-anchrored loops present at *Slc51b* locus (Fig. 4e). In contrast, the number of Foxa2-anchored loops drastically increases at a number of FXR targets with ligand-dependent FXR binding, including its own locus ^6^ (*Nr1i2*/PXR, *Abcb11*/Bsep, Cyp3a11, *Nr1h4*/FXR Fig. 4f).

Next, we identified Foxa2-anchored interchromosomal loops (1581 for vehicle, 8463 for GW4064, Fig. 5a) and they were mostly distinct for the two conditions (overlap 84 loops). Visualization of interchromosomal loops on chromosome 11 (vehicle, left panel; GW4064, middle panel; both, right panel) demonstrates a substantial increase in these interactions with addition of FXR agonist. We have reported that liver-specific *Foxa2* mutants accumulate hepatic bile acids due to decreased transcription of genes encoding bile acid transporters and bile acid conjugation enzymes (*Abcc2, Abcc3, Abcc4, Slco1a4, Slc27a5*) ^6,7^. We observe novel Foxa2-anchored interchromosomal loops forming in GW-treated livers for these targets responsible for cholestatic phenotype in *Foxa2*-deficient livers.

**Figure 5.**
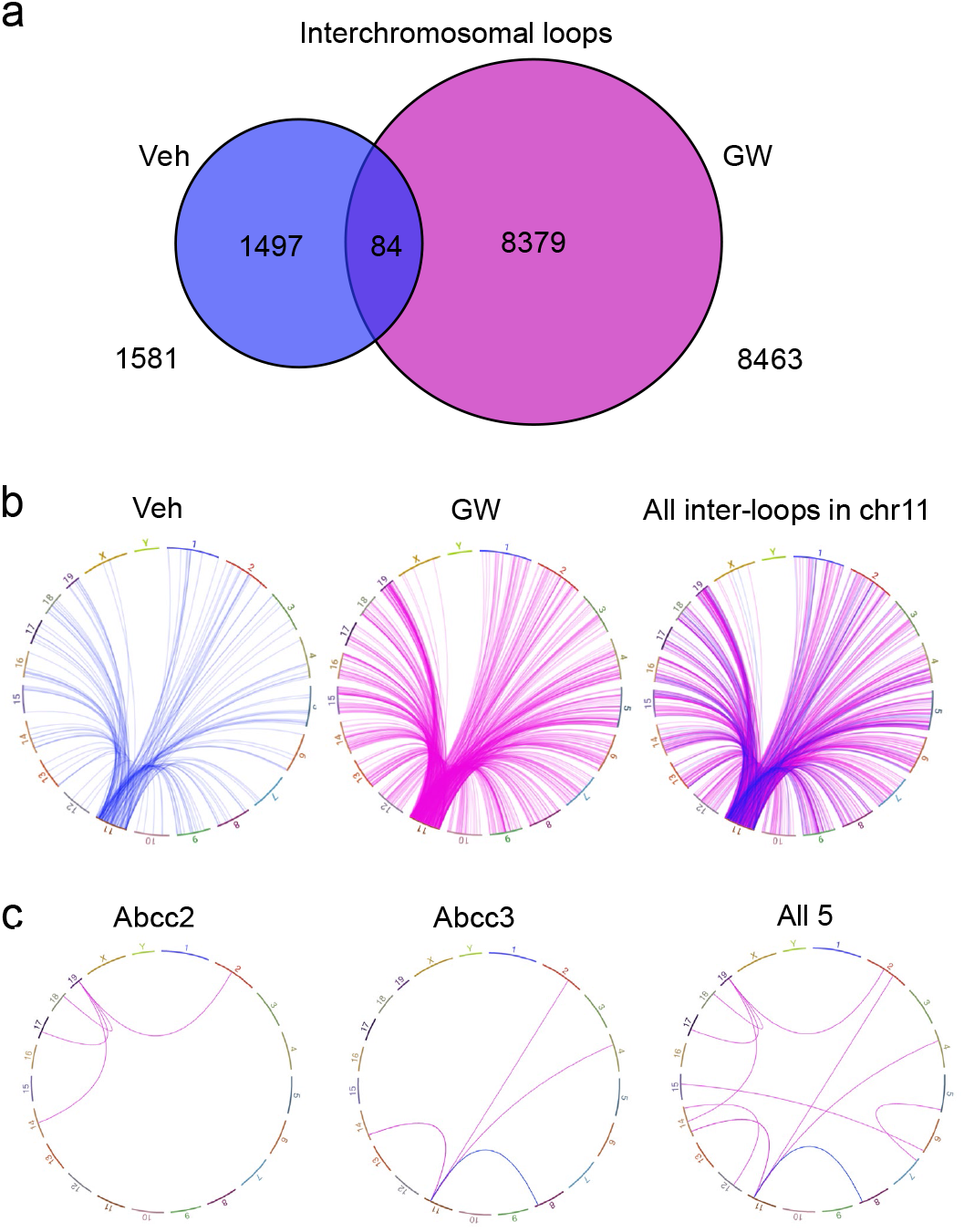
Foxa2-anchored interchromosomal loops are significantly increased in the presence of FXR ligand for activation. (**a**) Venn diagram comparing Foxa2-anchored interchromosomal loops in vehicle and GW4064-treated livers. (1581 for vehicle, 8463 for GW4064, 84 common) shows they were mostly distinct for the two conditions. (**b**) Visualization (IGV circle view) of interchromosomal loops on chromosome 11 (vehicle, left panel; GW4064, middle panel; both, right panel) demonstrates a substantial increase in these interactions with addition of FXR agonist. (**c**) Novel Foxa2-anchored interchromosomal loops form in GW-treated livers at Foxa2 targets responsible for cholestatic phenotype in *Foxa2*-deficient livers ^7^.

### Foxa2 and FXR ligand-dependent binding correlate with increase in intrachromosomal loops anchored by Foxa2 and activation of FXR targets

To ascertain whether chromatin conformation changes observed with GW treatment had functional consequences, we integrated interaction data with Foxa2 & FXR binding (Foxa2 ChIP-Seq, FXR ChIP-Seq) and differential gene expression in agonist treated livers (RNA-Seq) for FXR targets with ligand-dependent binding. We observe that increase in Foxa2-anchored intrachromosomal loops correlates with increased binding of both Foxa2 and FXR and activation of FXR-dependent gene transcription at these loci (*Abcb11*/Bsep, Cyp3a11, *Nr1h4*/FXR, *Nr1i2*/PXR, Fig.6).

**Figure 6.**
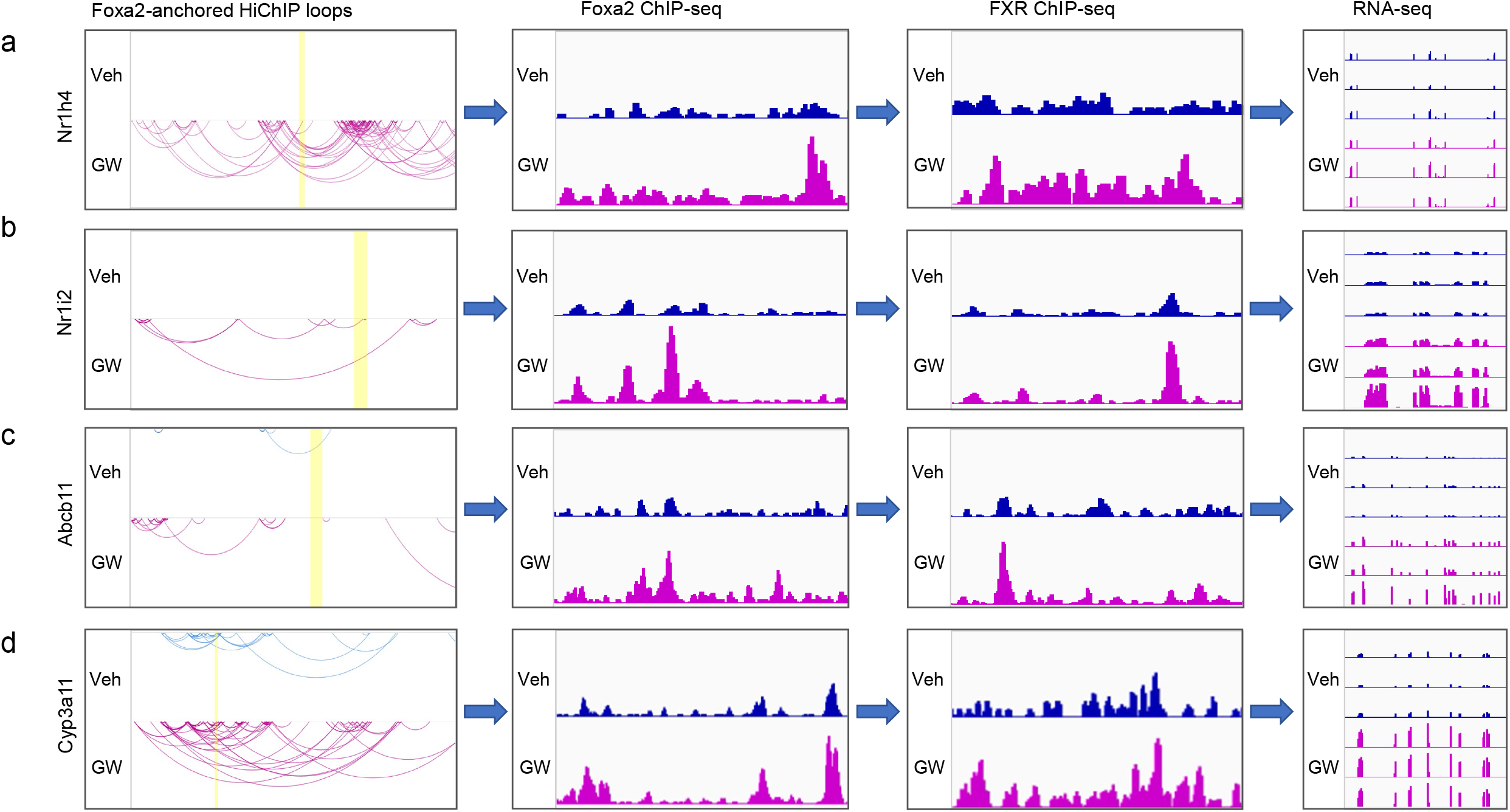
Foxa2 and FXR ligand-dependent binding correlate with increase in intrachromosomal loops anchored by Foxa2 and activation of FXR targets. Increase in Foxa2-anchored intrachromosomal loops correlates with increased binding of both Foxa2 and FXR and activation of FXR-dependent gene transcription at loci with FXR ligand-dependent binding (*Abcb11*/Bsep, Cyp3a11, *Nr1h4*/FXR, *Nr1i2*/PXR).

In summary, we demonstrate that genome-wide chromatin interactions are significantly expanded with ligand activation of FXR. Distribution of TADs changes and the number of Foxa2-anchored loops greatly increase with agonist treatment. Hence, we have identified a novel role for Foxa2 in chromatin conformation dynamics and extended its role in bile acid metabolism.

## DISCUSSION

We have previously challenged the accepted ligand-independent binding mechanism, showing that FXR and LXRα bind to both ligand-independent and ligand-dependent sites, inaccessible without the ligand, and Foxa2 modulates changes in chromatin accessibility for additional binding during ligand activation ^6^. In this study, we extend our findings, demonstrating drastic alterations in chromatin conformation accompany changes in chromatin accessibility with addition of agonist in ligand-dependent activation of FXR. These results also contest previously accepted mechanism of ligand-dependent activation involving ligand-independent binding. Since we demonstrated that ligand-dependent activation of both FXR and LXRα requires opening of chromatin in a common mechanism, we anticipate that chromatin conformation changes observed with addition of FXR agonist will also be shared in ligand-dependent activation of LXRα and likely for other type II nuclear receptors.

We demonstrate that the transformation of chromatin landscape with addition of FXR ligand is extensive, far more than expected. We observe a substantial increase in genome-wide interactions, reshuffling of TADs, and compartment switching in GW-treated livers. In addition, the number of both intrachromosomal and interchromosomal Foxa2-anchored loops dramatically changes with ligand addition. Our observations suggest that in addition to opening of chromatin for additional binding, there is a need for substantial chromatin restructuring and additional genome-wide interactions in order to achieve ligand-dependent activation of gene expression. These findings need to be taken into consideration in development of drug targets to treat medical conditions, such as metabolic disease and cancer.

We have described an extensive role for Foxa2 in hepatic bile acid metabolism ^6-8^ and synergistic cooperation between Foxa2 and bile acid receptor FXR in activating ligand-dependent gene expression ^6,8^. Here we demonstrate that Foxa2 also mediates chromatin conformation changes that accompany activation of FXR. Specifically, we observe drastic increase in Foxa2-anchored intrachromosomal loops at FXR locus and loci of FXR targets *Abcb11*/Bsep, Cyp3a11, and *Nr1i2*/PXR as well as novel Foxa2-anchored interchromosomal loops at Foxa2 targets that are responsible for cholestatic phenotype. Hence, Foxa2 plays a new role in enabling these structural changes, extending its function in bile acid homeostasis.

## ONLINE METHODS

### Mouse Liver Tissues

Wildtype male mice, 8–12 weeks of age, were used for all studies. Ligand activation was performed as described previously ^6^. Briefly, mice were treated once with GW4064, FXR agonist, using oral gavage. Control mice were treated with vehicle (20 mL propylene glycol/5 mL Tween 80 solution). Agonist was added to the vehicle for experimental treatment. Animals on both control and experimental treatments were sacrificed 4 h after gavage. All animal work was approved by Animal Care and Use Committee at UVa (protocol number 4162–03–20).

### HiChIP Assay

HiChIP experiments were performed with vehicle- and ligand-treated mouse liver tissues using the Arima-HiC+ Kit (Arima Genomics, Cat. A101020) according to its recommended protocol.

#### Pulverization and Crosslinking of Liver Tissues

About ∽200 mg of each mouse liver tissue was pulverized using Covaris CryoPrep System with impact level setting 3. Pulverized tissues were washed with cold 1xPBS and pelleted by centrifugation at 1,000xg at 4°C for 5 min. 3ml of TLB1 was added to resuspend each sample. After incubation for 20 min at 4°C, samples were filtered through 40 μm cell strainers. The flowthrough was pelleted by centrifugation at 1,000×g at 4°C for 5 min and resuspended in 1 ml of TLB2. Then 3 ml of Sucrose Solution (SS) was carefully overlaid on top of the samples and pelleted by centrifugation at 4°C for 5 min. Samples were then resuspended in 5 ml 1XPBS and crosslinked by adding 286 μl of 37% formaldehyde to achieve a final concentration of 2% (wt/vol) and incubated at room temperature for 10 min. Then 460 μl of Stop Solution 1 was added, samples were incubated at room temperature for 5 min, and then on ice for 15 min. Samples were pelleted by centrifugation at 2,500 × g at 4°C for 5 min and resuspended in 1 ml cold 1xPBS. An aliquot which contains the equivalent of 10 mg of original pulverized tissue for each sample was then measured by Qubit 4 Fluorometer for total DNA yield.

#### HiChIP Preparation

The crosslinked samples which contained the equivalent of 15 μg of total DNA for each sample were resuspended in 20 μl of lysis buffer and incubated at 4°C for 20 min. Then 24 μl of Conditioning Solution was added and samples were incubated at 62 °C for 10 min. Then 20 μl of Stop Solution 2 was added and samples were placed at 37°C for 15 min. A master mix of 7 μl of buffer A, 1 μl of enzyme1 μl of restriction enzyme MboI and 1 μl of restriction enzyme of HinfI were added to each sample, and samples were incubated at 37°C for 1 hr. Samples were then pelleted at 2,500 × g at 4°C for 5 min. Next, samples were gently rinsed in 1.5 ml of dH_2_O, pelleted, and resuspended in 75 μl of dH_2_O. Then 16 μl of master mix containing 12 μl of Buffer B and 4 μl of Enzyme B was added to each sample, mixed well and incubated at room temperature for 45 min. Next, 82 μl master mix containing 70 μl of Buffer C and 12 μl of Enzyme C was added to each sample, mixed well and incubated at room temperature for 15 min. Samples were then pelleted and resuspended in 110 μl of cold R1 Buffer (10 mM Tris-HCl pH8.0, 140 mM NaCl, 1 mM EDTA, 1% Triton X-100, 0.1% SDS, 0.1% sodium deoxycholate) and incubated at 4°C for 20 min. Chromatin shearing was then performed using Diagenode Bioruptor Pico with “30sec ON/30sec OFF” condition for 11-12 cycles. Then 900 μl of R1 Buffer was added to each sheared sample to get total volume of 1 ml. For Foxa2 HiChIP immunoprecipitation, 3.5 μl of Foxa2 antibody (Seven Hills Bioreagents, Cat: WRAB-1200) was added to each sample and incubated at 4°C overnight with rotating. Next day, 30 μl of pre-blocked Protein G beads was added to each sample and incubated at 4°C for 4 hr with rotating. After incubation, beads were sequentially washed three times with R1 Buffer, two times with R3 Buffer (0.1% SDS, 1% Triton X-100, 2 mM EDTA, 20 mM Tris–HCl pH 8.5, 0.1% sodium deoxycholate and 300 mM NaCl), one time with LC Buffer (1 mM EDTA, 10 mM Tris–HCl pH 8.5, 0.1% sodium deoxycholate and 150 mM lithium chloride, 0.5% IGEPAL CO-630), and two times with LTE Buffer (10 mM Tris–HCl pH 8.0, and 0.1 mM EDTA). The beads of each sample were resuspended with 174 μl of Elution Buffer. Then 35.5 μl of master mix containing 10.5 μl of Buffer D and 25 μl of Enzyme D and 20 μl of Buffer E was added to each sample, and incubated at 55°C for 30 min, 68°C for 90 min, and 25°C for 10 min. The bead-bound DNA for each sample was purified with 230 μl of AMpure XP beads (Beckman Coulter, Cat# A638880) and washed two times with 700 μl of 80% ethanol, then resuspended thoroughly in 50 μl of Elution Buffer. DNA concentrations were measured using Qubit 1x dsDNA HS Assay Kit (Invitrogen by ThermoFisher Scientific, Cat: Q33231).

#### Biotin Enrichment and HiChIP Library Preparation

Elution Buffer was added to beads-bound DNA to bring the total volume to 100 μl for each sample. Then 100 μl of Enrichment Beads were added, mixed well and incubated at RT for 15 min. Beads were washed two times with 200 μl of Wash Buffer and one time with 100 μl of Elution Buffer, then resuspended in 40 μl of Elution Buffer.

For end-repair and adaptor ligation, 20 μl of a master mix containing 13 μl of Low EDTA TE, 6 μl of Buffer W1 and 1 μl of Enzyme W2 was added to each sample, and incubated at 37°C for 10 min. Beads were washed twice with 150 μl of Wash Buffer and once with 100 μl of Elution Buffer, then resuspended in 50 μl of a master mix containing 30 μl of Low EDTA TE, 5 μl of Buffer G1, 13 μl of Reagent G2, 1 μl of Enzyme G3 and 1 μl Enzyme G4, and incubated at 20°C for 20min. Beads were washed twice with 150 μl of Wash Buffer and once with 100 μl of Elution Buffer. Then beads were resuspended in 25 μl of a master mix containing 20 μl of Low EDTA TE, 3 μl of Buffer Y, 2 μl of Enzyme Y3 and 5 μl of uniquely indexed Reagent Y2, mixed well and incubated at 25°C for 15 min. Beads were washed twice with 150 μl of Wash Buffer and once with 100 μl of Elution Buffer. Then resuspended in 50 μl of a master mix containing 30 μl of Low EDTA TE, 5 μl of Buffer B1, 2 μl of Reagent B2, 9 μl of Reagent B3, 1 μl of Enzyme B4, 2 μl of Enzyme B5 and 1 μl of Enzyme B6, mixed well and incubated at 40°C for 10 min. Beads were washed twice with 150 μl of Wash Buffer and one time with 100 μl of Elution Buffer. Then resuspended in 22 μl of Elution Buffer. These HiChIP libraries were then amplified using Accel-NGS 2S Plus DNA Library Kit for Illumina Platforms (Swift Biosciences, Cat# 21024) and Accel-NGS 2S Indexing Kit (Set A) for illumine Platforms (Swift Biosciences, Cat# 26148).

#### Illumina Sequencing

The HiChIP libraries were deep sequenced using the Next-Seq Sequencer 2000 (Illumina) in paired-end 2 × 150 bp mode using the NextSeq™ 1000/2000 P2 Reagents (300 Cycles) sequencing kit (Illumina, Cat# 20044466).

### HiChIP Data Analysis

#### HiChIP Data Processing with HiC-Pro

HiChIP paired-end reads were aligned to mm10 genome assembly using the HiC-Pro pipeline ^10^ using bowtie2 (version 2.2.9) and with mapping quality filter 15. Default settings were used to remove singleton, multi-mapped and duplicate reads. Reads were assigned to MboI (*^GATC*) and HinfI (*G^ANTC*) restriction fragments and filtered for valid interactions using min and max size parameters of 100 and 100,000, and binned interaction matrices were generated.

#### HiChIP Significant Contact Calling with HiCCUPS and Fit-HiC

HiCCUPS ^19^ did not run on the interaction matrix because of sparsity. Instead, we used Fit-Hi-C contact caller ^20^. Bin pairs of the interaction matrix with statistically significant contact signals were identified using Fit-Hi-C. Genome wide intra-chromosomal bin pairs were filtered for an interaction distance between 20kb and 2Mb, and default Fit-HiC settings were used to calculate false discovery rate (FDR) values for each bin-pair in a given HiChIP experiment. Bin size of 50kb and FDR cutoff < 0.001 were used to identify significant contacts.

#### HiChIP Differential Significant Contact Calling with HiCcompare

HiCcompare ^21^ uses the raw Hi-C matrix of HiC-Pro output as input. Two Hi-C matrices were jointly normalized for each chromosome. The default settings were used. The interactions with low average expression were filtered out with A.min=15. The value of FDR<0.05 was used for differential contact selection.

#### Topological Associating Domain (TAD) Calling with HiCExplorer

TADs were called with HiCExplorer ^22^. Hi-C contact matrix was used as input. TAD-separation scores were calculated for all bins. The TAD-separation scores were evaluated and the cutoff p-values < 0.05 were applied to select the TAD boundaries.

#### Compartment A/B Profile Calling with cworld

Compartment was called using cworld ^23^. Genomic bins of reference genome were generated at 250kb. The command matrix2compartment command was used to extract the compartment A/B profiles with default settings.

#### Foxa2-Anchored Loops Calling with hichipper

The hichipper (version 0.7.7) ^24^ was used to identify the Foxa2-anchored chromatin loops. The HiC-Pro matrix output, restriction digested mm10 genome assembly and ChIP-Seq Foxa2 binding sites ^6^ were used as input to identify chromatin loops extending from regions occupied by Foxa2. The default parameter settings were used with minimum distance of 5000 and maximum peak padding width of 2000000.

#### Visualization of HiChIP Matrices, Loops and Compartment A/B Profiles

Hi-C matrices were visualized using Juicebox ^17^. Foxa2-anchored intra- and inter-chromosomal loops were visualized using WashU Epigenome Browser ^25^, Integrative Genome Viewer (IGV)^18^ and UCSC Genome Browser. Compartment A/B profiles were visualized using UCSC Genome Browser ^26^.

#### Functional Analysis

Interaction regions were associated with closest genes using GREAT ^11^, which were subsequently used for pathway analysis with Enrichr ^12^ and Ingenuity Pathway Analysis as described previously ^6,27^. Scanning motif analysis for overrepresented transcription factor binding motifs in interaction regions was performed by PscanChIP ^28^. Heatmaps of HiChIP coverage were generated by deepTools ^29^. The overlap between different categories of interactions was computed using bedtools (version 2.29.2) ^30^.

### Data Availability

Genomic data from this study (Foxa2 HiChIP in vehicle and GW4064-treated livers) can be accessed at GEO accession number XXX Foxa2 ChIP-Seq, FXR ChIP-Seq, and RNA-Seq data for vehicle- and GW4064-treated livers can be found at GEO accession number GSE149075.

## EXTENDED DATA FIGURE LEGENDS

**Figure 1.**
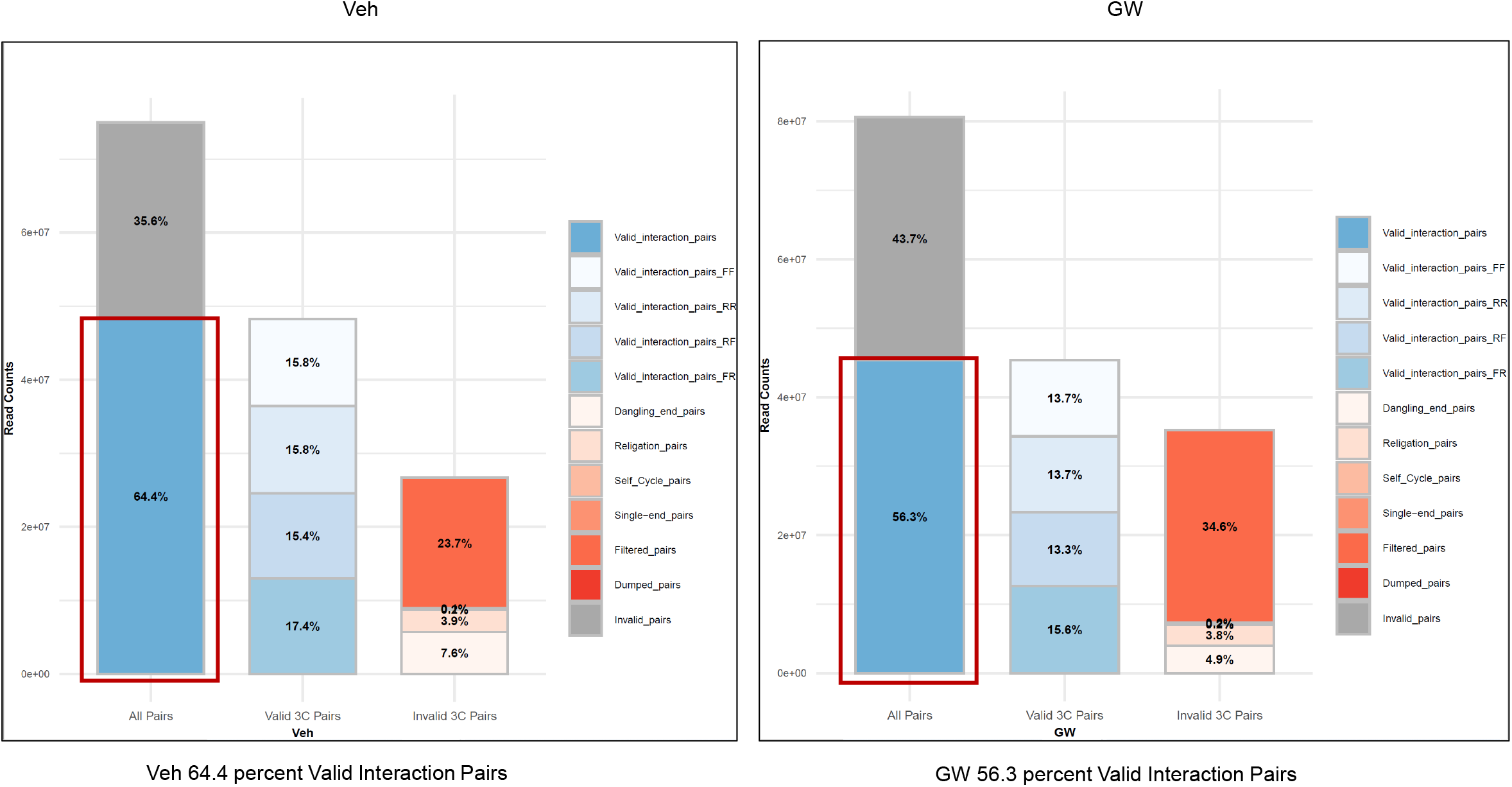
HiChIP Library alignment statistics. Quality control metrics from HiC-Pro showing percentage of valid pairs in HiChIP libraries (64.4% vehicle, 56.3 % GW4064) that were used for subsequent analyses.

**Figure 2.**
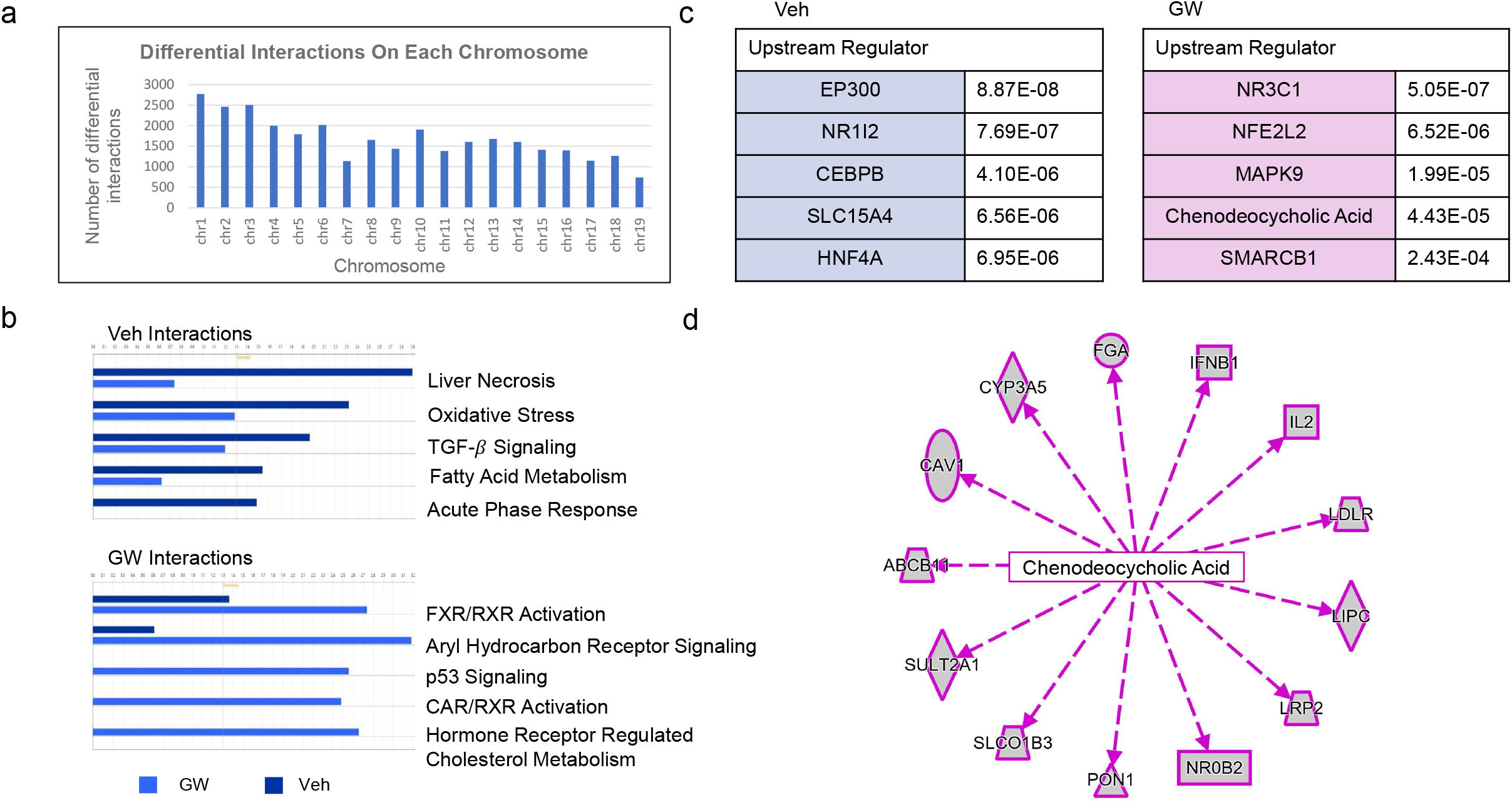
Functional analysis of genome-wide differential interactions. **(a)** A plot showing the number of differential interactions on each chromosome.(**b**) Pathways identified by Ingenuity pathway analysis (IPA) of genes in differential interactions regions include “ Oxidative Stress”, “Fatty Acid Metabolism” and “Acute Phase Response” in control livers while genes in FXR and CAR activation pathways and those regulating cholesterol metabolism were found in agonist-treated livers, consistent with bile acid activation. (**c**)IPA of genes in differential interaction regions identified multiple upstream regulators, including HNF4α, CEBPB, and EP300 for control livers and NFE2L2, SMARCB1, and chenodeoxycholic acid. (**d**) Chenodeoxycholic acid-regulated network identified by IPA for GW-treated livers. Chenodeoxycholic acid is a naturally occurring bile acid that activates FXR. Hence, IPA analysis confirms FXR activation in agonist-treated livers.

**Figure 3.**
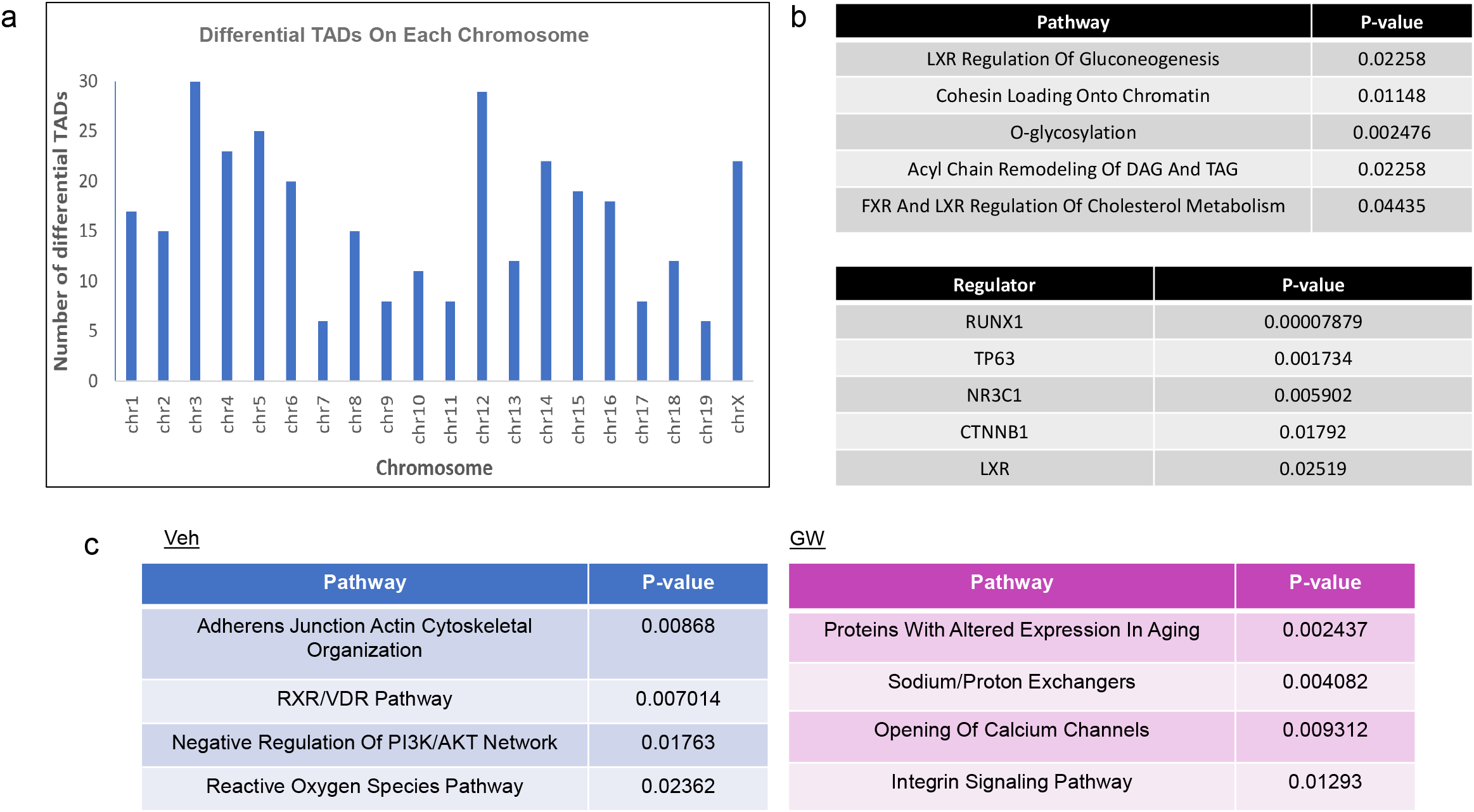
Functional analysis of topological associated domains (TADs) **(a)** A plot showing the number of differential TADs on each chromosome. (**b**) Enrichr pathway analysis of genes in differential TAD regions identified overrepresented pathways including “LXR regulation of gluconeogenesis” and “FXR and LXR regulation of cholesterol metabolism”, consistent with bile acid activation, as well as “Cohesin loading onto chromatin”, congruous with cohesin function in mediating chromatin loop formation (top panel). EnrichR ChEA analysis of genes in differential TADs identified overlap with RUNX1, TP53, β-catenin, and LXR targets (bottom panel). (**c**) Functional analysis of genes in switching compartments shows “Cytoskeletal Organization”, “Reactive Oxygen Species”, and “RXR/VDR pathway” pathways overrepresented in 54 regions repressed with agonist treatment and “Proteins with Altered Expression in Aging”, “Sodium/Proton Exchangers”, and “Opening of Calcium Channels” in 152 regions activated with GW treatment.

**Figure 4.**
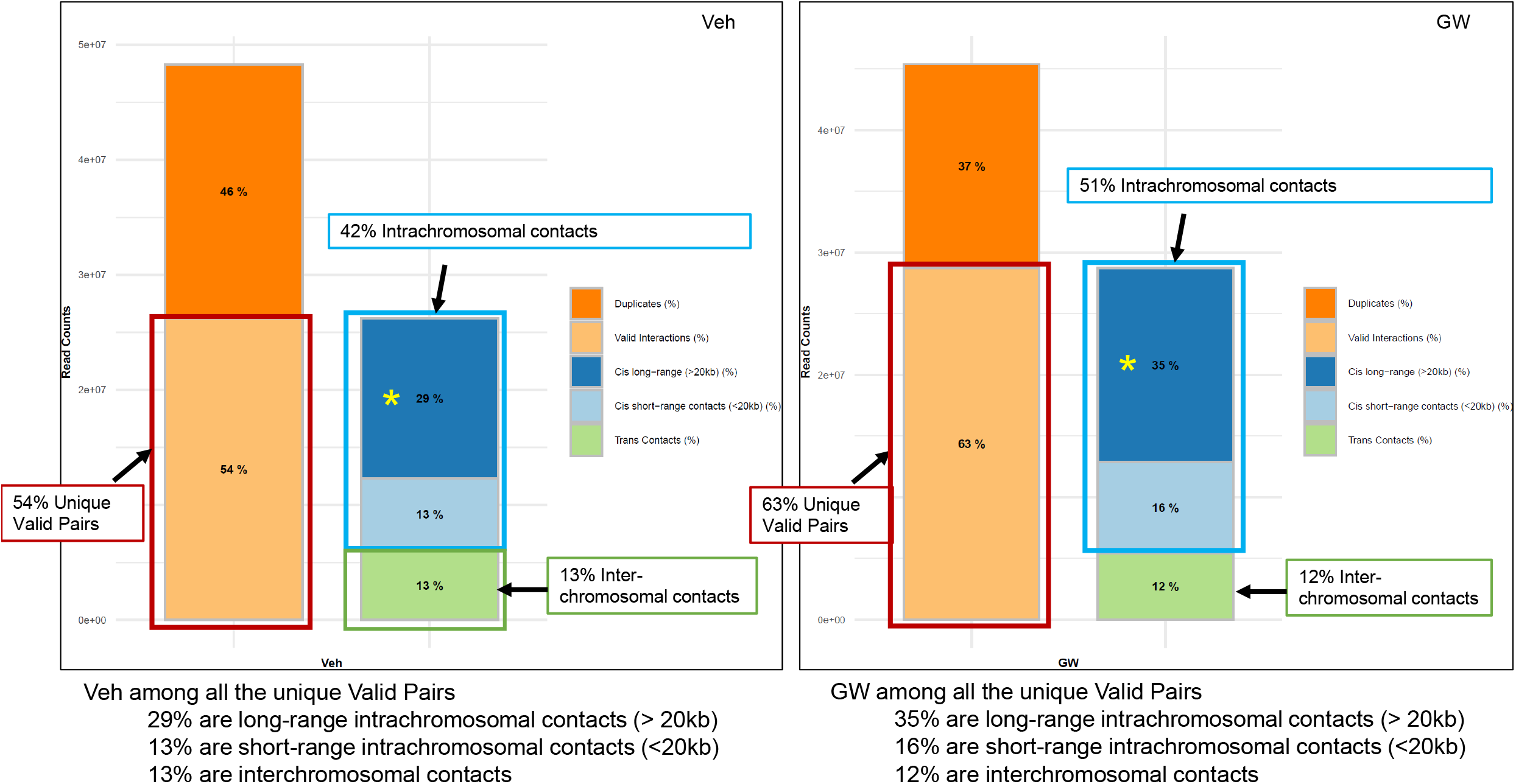
Statistics of intrachromosomal and interchromosomal contacts of HiChIP libraries. Quality control metrics from HiC-Pro showing percentage of unique valid pairs (duplicates removed) in HiChIP libraries with intrachromosomal and interchromosomal contacts.

## COMPETING INTERESTS

The authors have nothing to declare.

## ACKNOWLEDGEMENTS

We thank J.Schinderle, N.Reddy, and B. Weidemann for technical assistance. I.M.B. is supported by National Institute of Diabetes and Digestive and Kidney Diseases R01 award DK121059

## AUTHOR CONTRIBUTIONS

**YH**: Investigation, Formal Analysis, Software, Visualization, Writing - Review & Editing; **LH**: Software, Formal Analysis, Data Curation, Visualization, Writing - Review & Editing; **AW:** Software, Formal Analysis, Data Curation, Visualization, Writing - Review & Editing;; **IMB:** Conceptualization, Formal Analysis, Investigation, Writing – original draft, Supervision, Project administration, Funding acquisition.

## Notes

### Competing Interest Statement

The authors have declared no competing interest.

## REFERENCES

1. Lin, C.Y. & Gustafsson, J.A. Targeting liver X receptors in cancer therapeutics. Nat Rev Cancer 15, 216–24 (2015).

2. Wright, M.B., Bortolini, M., Tadayyon, M. & Bopst, M. Minireview: Challenges and opportunities in development of PPAR agonists. Mol Endocrinol 28, 1756–68 (2014).

3. Schaap, F.G., Trauner, M. & Jansen, P.L. Bile acid receptors as targets for drug development. Nat Rev Gastroenterol Hepatol 11, 55–67 (2014).

4. Ma, Z. et al. Liver X Receptors and their Agonists: Targeting for Cholesterol Homeostasis and Cardiovascular Diseases. Curr Issues Mol Biol 22, 41–64 (2017).

5. Kidani, Y. & Bensinger, S.J. Liver X receptor and peroxisome proliferator-activated receptor as integrators of lipid homeostasis and immunity. Immunol Rev 249, 72–83 (2012).

6. Kain, J. et al. Pioneer factor Foxa2 enables ligand-dependent activation of type II nuclear receptors FXR and LXRα. Mol Metab 53, 101291 (2021).

7. Bochkis, I.M. et al. Hepatocyte-specific ablation of Foxa2 alters bile acid homeostasis and results in endoplasmic reticulum stress. Nat Med 14, 828–36 (2008).

8. Bochkis, I.M. et al. Foxa2-dependent hepatic gene regulatory networks depend on physiological state. Physiol Genomics 38, 186–95 (2009).

9. Bochkis, I.M., Shin, S. & Kaestner, K.H. Bile acid-induced inflammatory signaling in mice lacking Foxa2 in the liver leads to activation of mTOR and age-onset obesity. Molecular Metabolism 2, 447–456 (2013).

10. Servant, N. et al. HiC-Pro: an optimized and flexible pipeline for Hi-C data processing. Genome Biology 16, 259 (2015).

11. McLean, C.Y. et al. GREAT improves functional interpretation of cis-regulatory regions. Nat Biotechnol 28, 495–501 (2010).

12. Kuleshov, M.V. et al. Enrichr: a comprehensive gene set enrichment analysis web server 2016 update. Nucleic Acids Res 44, W90–7 (2016).

13. Chien, R., Zeng, W., Ball, A.R. & Yokomori, K. Cohesin: a critical chromatin organizer in mammalian gene regulation. Biochem Cell Biol 89, 445–58 (2011).

14. Zhang, L.A.-O. et al. Runt-related transcription factor-1 ameliorates bile acid-induced hepatic inflammation in cholestasis through JAK/STAT3 signaling. LID - 10.1097/HEP.0000000000000041 [doi].

15. Kim, D.H. & Lee, J.W. Tumor suppressor p53 regulates bile acid homeostasis via small heterodimer partner. Proc Natl Acad Sci U S A 108, 12266–70 (2011).

16. Thompson, M.D. et al. β-Catenin regulation of farnesoid X receptor signaling and bile acid metabolism during murine cholestasis. Hepatology 67, 955–971 (2018).

17. Durand, N.C. et al. Juicebox Provides a Visualization System for Hi-C Contact Maps with Unlimited Zoom. Cell Syst 3, 99–101 (2016).

18. Robinson, J.T. et al. Integrative genomics viewer. Nat Biotechnol 29, 24–6 (2011).

19. Rao, S.S. et al. A 3D map of the human genome at kilobase resolution reveals principles of chromatin looping. Cell 159, 1665–80 (2014).

20. Ay, F., Bailey, T.L. & Noble, W.S. Statistical confidence estimation for Hi-C data reveals regulatory chromatin contacts. Genome Res 24, 999–1011 (2014).

21. Stansfield, J.C., Cresswell, K.G., Vladimirov, V.I. & Dozmorov, M.G. HiCcompare: an R-package for joint normalization and comparison of HI-C datasets. BMC Bioinformatics 19, 279 (2018).

22. Ramírez, F. et al. High-resolution TADs reveal DNA sequences underlying genome organization in flies. Nature Communications 9, 189 (2018).

23. Sanders, J.T. et al. Loops, topologically associating domains, compartments, and territories are elastic and robust to dramatic nuclear volume swelling. Scientific Reports 12, 4721 (2022).

24. Lareau, C.A. & Aryee, M.J. hichipper: a preprocessing pipeline for calling DNA loops from HiChIP data. Nature Methods 15, 155–156 (2018).

25. Li, D. et al. WashU Epigenome Browser update 2022. Nucleic Acids Res 50, W774–81 (2022).

26. Kent, W.J. et al. The human genome browser at UCSC. Genome Res 12, 996–1006 (2002).

27. Wei, X. et al. Redistribution of lamina-associated domains reshapes binding of pioneer factor FOXA2 in development of nonalcoholic fatty liver disease. Genome Res 32, 1981–1992 (2022).

28. Zambelli, F., Pesole, G. & Pavesi, G. PscanChIP: Finding over-represented transcription factor-binding site motifs and their correlations in sequences from ChIP-Seq experiments. Nucleic Acids Res 41, W535–43 (2013).

29. Ramirez, F., Dundar, F., Diehl, S., Gruning, B.A. & Manke, T. deepTools: a flexible platform for exploring deep-sequencing data. Nucleic Acids Res 42, W187–91 (2014).

30. Quinlan, A.R. & Hall, I.M. BEDTools: a flexible suite of utilities for comparing genomic features. Bioinformatics 26, 841–2 (2010).

